# Unearthing SRSF1’s Novel Function in Binding and Unfolding of RNA G-Quadruplexes

**DOI:** 10.1101/2023.10.30.563137

**Authors:** Naiduwadura Ivon Upekala De Silva, Nathan Lehman, Talia Fargason, Trenton Paul, Zihan Zhang, Jun Zhang

## Abstract

SRSF1 governs splicing of over 1,500 mRNA transcripts. SRSF1 contains two RNA-recognition motifs (RRMs) and a C-terminal Arg/Ser-rich region (RS). It has been thought that SRSF1 RRMs exclusively recognize single-stranded exonic splicing enhancers, while RS lacks RNA-binding specificity. With our success in solving the insolubility problem of SRSF1, we can explore the unknown RNA-binding landscape of SRSF1. We find that SRSF1 RS prefers purine over pyrimidine. Moreover, SRSF1 binds to the G-quadruplex (GQ) from the ARPC2 mRNA, with both RRMs and RS being crucial. Our binding assays show that the traditional RNA-binding sites on the RRM tandem and the Arg in RS are responsible for GQ binding. Interestingly, our FRET and circular dichroism data reveal that SRSF1 unfolds the ARPC2 GQ, with RS leading unfolding and RRMs aiding. Our saturation transfer difference NMR results discover that Arg residues in SRSF1 RS interact with the guanine base but other nucleobases, underscoring the uniqueness of the Arg/guanine interaction. Our luciferase assays confirm that SRSF1 can alleviate the inhibitory effect of GQ on gene expression in the cell. Given the prevalence of RNA GQ and SR proteins, our findings unveil unexplored SR protein functions with broad implications in RNA splicing and translation.

## Introduction

RNA metabolism is fundamental to cell differentiation and tissue development. Tight regulation of RNA metabolism is orchestrated by numerous RNA-binding proteins and dynamic RNA secondary structures(1–3). Ser/Arg-rich (SR) proteins are a family of RNA-binding proteins that regulate alternative splicing(4–8), a process that governs 95% of human genes, enabling 20,000 protein-encoding genes to produce 200,000 protein isoforms(9–12). The SR protein family consists of 12 members (SRSF1-SRSF12), each featuring one to two N-terminal RNA recognition motifs (RRMs) and a phosphorylatable C-terminal protein region rich in repetitive Arg/Ser dipeptides (RS) (13,14). SR proteins typically recognize exonic splicing enhancers and promote inclusion of bound exons, although recent studies have found that the effects of SR proteins depend on where they bind. For example, binding of SR proteins to introns inhibits the use of the neighboring splicing site for some RNA transcripts (15). In addition to their roles in regulating RNA splicing, SR proteins also affect genome stability(16), mRNA transcription(17,18), mRNA transport(19), translation(20,21), nonsense-mediated mRNA decay(22), and regulation of long noncoding RNA(23,24).

The RRM domains of SR proteins exhibit distinct but broad RNA-binding specificities, believed to play a dominant role in determining the RNA motifs that SR proteins recognize. For instance, SRSF1 contains two RRMs (RRM1 and RRM2, in a tandem configuration). RRM1 prefers C-containing RNA motifs (25), while RRM2 prefers GGA motifs (26,27) (Fig. 1A). Consequently, SRSF1’s RRM tandem recognizes purine-rich RNA motifs with upstream or downstream C motifs (25,28). Unlike SRSF1, SRSF3’s RRM recognizes pyrimidine-rich RNA sequences(29). Due to their broad specificity, each SR protein can govern processing of many mRNA transcripts. For instance, SRSF1 alone affects the splicing of over 1,500 RNA transcripts(30).

**Figure 1:**
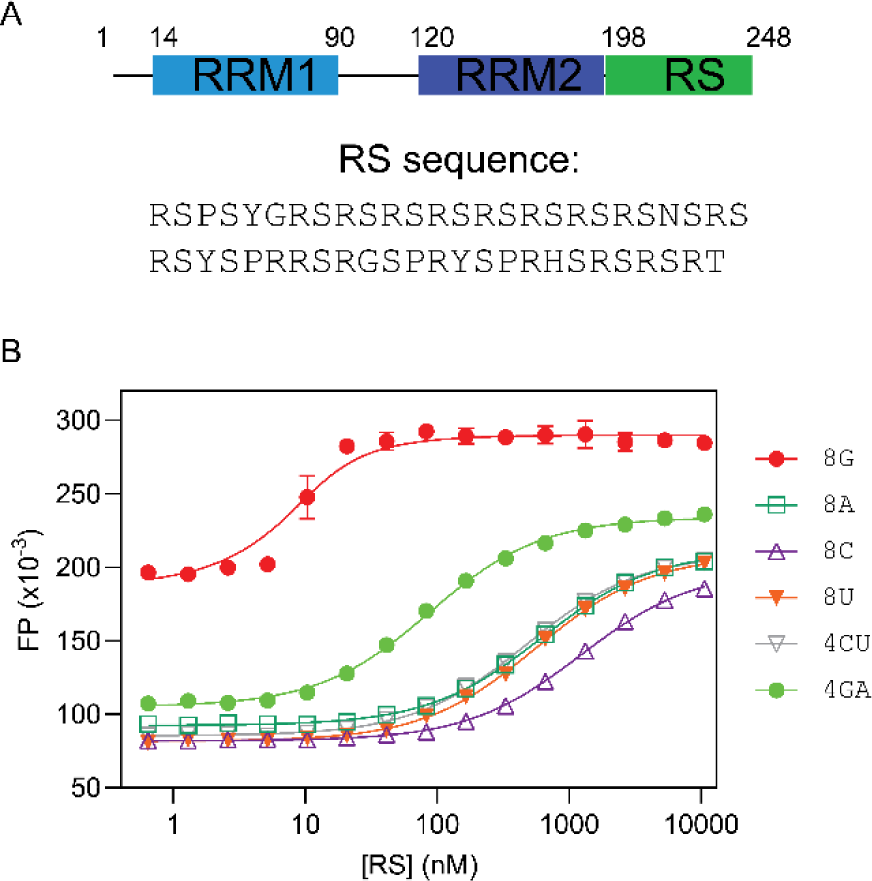
SRSF1 RS prefers purine ribonucleotides. (A) Domain architecture of SRSF1. The residue numbers are shown for these domain boundaries. The RS sequence (residues 198-248) are shown blow. (B) Fluorescence polarization binding profiles for RS with various 8-mer RNA nucleotides. RNA oligomers were labeled by fluorescein at the 5’ end, and fluorescent probe concentration was 10 nM for all binding assays. The binding assays were performed in a buffer containing 35 mM K_3_PO_4_, 2.5 mM NaCl, 15 mM HEPES pH 7.5, 0.015% TWEEN 20, 0.075 mM TCEP.

RS domains within SR proteins exhibit diverse lengths, ranging from 50 amino acids in SRSF1 to around 300 amino acids in SRSF4. Most of RS dipeptide repeats in SR proteins undergo varying degrees of phosphorylation on serine residues. For example, the RS domain of SRSF1 can exist in an unphosphorylated state, be partially phosphorylated (with the addition of 12 phosphate groups), or become hyperphosphorylated (with the addition of 18 phosphate groups). These RS domains are widely thought to interact with other protein factors that participate in splicing, such as U1 and U2 particles (31–41). Furthermore, these interactions are regulated by phosphorylation or dephosphorylation, which are pivotal in coordinating the spliceosome assembly and activation (31–41). Notably, RS domains of SR proteins potentially engage in RNA binding. A UV crosslinking experiment has shown a direct interaction of RS with RNA (42,43). Studies on IgM splicing have highlighted the indispensability of RS domains, as the exonic splicing enhancer presents sub-optimal binding sites for RRMs (44–46). RS domains are also present in a larger family called SR-like proteins, which are extensively involved in RNA processing(47–50). These observations hint at the potential role of RS domains as crucial RNA-binding components.

RNA metabolism is also subject to regulation by RNA structure, with one RNA structure garnering increasing attention: G-quadruplexes (GQs). In a GQ, four guanine bases form a planar array known as a G-tetrad via Hoogsteen base pairing. Typically, three or more layers of G-tetrads stack to create a GQ structure. Mono cations reside between G-tetrad layers, bonding with carbonyl oxygen atoms of guanines, and their stabilization of GQ follows an order of K^+^ > Na^+^ > Li^+^. GQ structures can arise from both DNA and RNA sequences, and accumulating evidence underscores their roles in governing RNA transcription, stability, and RNA translation. Recent genome-wide exploration unveiled the presence of tens of thousands of GQs within the human transcriptome, with higher enrichment near exon-intron boundaries, implicating their involvement in alternative splicing(51). Despite the abundance of GQ-promoting sequence motifs, only a fraction of G-rich sequences yield functional GQ structures within cells(52). Remarkably, certain GQ formations are exceptionally stable, necessitating mechanisms to curtail extensive GQ assembly(53). Several RNA-binding proteins participate in this regulation. For example, hnRNPH1 has demonstrated its ability to inhibit GQ formation(54). Nevertheless, hnRNPH1’s RNA recognition is limited to a small subset of RNA molecules. The identity of other proteins that share this regulatory role remains unknown. Recent investigations have uncovered that SR proteins, among numerous other proteins, can be recruited by a GQ-forming RNA from actin related protein complex subunit 2 (ARPC2), which regulates actin polymerization and cell migration (55). The ARPC2 GQ is in the 5’ untranslated region of the ARPC2 mRNA. However, questions regarding the specificity of binding, the essential protein domains for GQ interaction, and the implications of this binding on ARPC2’s GQ structure are still unanswered.

With our recent success in obtaining soluble full-length SRSF1, we use multiple biophysical methods to characterize the RNA-binding specificity of the RS domain of SRSF1 and investigate the interaction between SRSF1 and the ARPC2 GQ. Contrary to the prevalent belief that the RS domain has no RNA-binding specificity, we find that the SRSF1 RS prefers purine-rich RNA. We show that SRSF1 binds with ARPC2 GQ with a nanomolar affinity, and both the RRM tandem and RS are needed for GQ binding. Using fluorescence resonance energy transfer (FRET), we find that SRSF1 unfolds the ARPC2 GQ. This finding is further confirmed by circular dichroism. Interestingly, the RS domain plays a dominant role in unfolding the ARPC2 GQ. Through mutagenesis, we found that arginine residues in RS are responsible for the unfolding of GQ. We further use saturation transfer difference (STD) NMR to show that arginine sidechains interact with guanine bases but not with other nucleotide bases, providing an atomic-level mechanism for GQ unfolding by RS domains. Our luciferase assays suggest that the ARPC2 GQ inhibits the Renilla luciferase expression, which is consistent with the typical effect of GQ on gene expression. Interestingly, this inhibition can be rescued by co-expression of full-length SRSF1 but the RRM tandem, suggesting the crucial role of RS. In summary, our study reveals a new function of SR proteins in binding and unfolding GQ RNA, indicating a broad impact in regulating RNA metabolism given the prevalence of GQ and SR proteins.

## Materials and Methods

### Synthetic RNA oligonucleotides

Synthetic 5’ fluorescein-labeled RNAs: ARPC2, 5’-AGCCGGGGGCUGGGCGGGGACCGGGCUUGU-3’; ARPC2, 5’-GGGGGCUGGGCGGGGACCGGG-3’; ARPC2 GQ mutant, 5’-GUAGACUGAGCGAAGACCGAG-3’; 21-mer GA, 5’-GAGAGAGAGAGAGAGAGAGAG-3’; 21-mer CU, 5’-CUCUCUCUCUCUCUCUCUCUC-3’; 5’-UUCAGAGGA-3’; 5’-UCAGAGGG-3’; 8G, 5’-GGGGGGGG-3’; 8C, 5’-CCCCCCCC-3’; 8U, 5’-UUUUUUUU-3’; 8A, 5’-AAAAAAAA-3’; 4GA, 5’-GAGAGAGA-3’; 4CU, 5’-CUCUCUCU-3’.

Synthetic Cy3-Cy5 or Cy3 labeled RNA: Cy3-Cy5 ARPC2 GQ, 5’-Cy5-UGGGGGCUGGGCGGGGACCGGGU-Cy3-3’; Cy3-ARPC2 GQ, 5’-UGGGGGCUGGGCGGGGACCGGGU-Cy3-3’

All RNA sequences, except for Cy3-Cy5 labeled RNA, were purchased from Dharmacon and dissolved in H_2_O to achieve a concentration of 400 μM. G-quadruplex formation of all 21-mer RNA and ARPC2 RNA (200 μM) was carried out in the folding buffer containing 20 mM Tris-HCl, pH 7.5, 25 mM KCl, and 0.1 mM EDTA. RNA was folded using a thermocycler by heating the RNA at 95^◦^C for 5 min, followed by an annealing gradient to 4^◦^C at a rate of 1 ^◦^C per minute. Folded RNA used in FP assays, CD, and UV melting experiments were diluted in 25 mM KCl, 0.25 M Arg/Glu, 20 mM Tris-HCl pH 7.5, 0.1 mM EDTA, 0.1 mM TCEP, 0.02 % TWEEN 20, to achieve the concentration required for each experiment. Cy3-Cy5 or Cy3-labeled RNA (Integrated DNA Technology, IDT) was first dissolved in H_2_O to achieve a concentration of 100 μM, and folded following the same procedure as stated above.

### Molecular cloning and protein expression

The human SRSF1 gene (residues 1−248) was cloned into pSMT3 using BamH I and Hind III. The RRM tandem (residues 1−196), the RS domain (residues199-248), and their mutants were prepared using mutagenesis PCR. All SRSF1 constructs were expressed by BL21-CodonPlus (DE3) cells cultured in LB media at 37 ^◦^C. Once the cell density reached an OD_600_ of 0.6, 0.5 mM of isopropyl thio-β-galactoside (IPTG) was added to induce protein expression at 22 °C for 16 hours. The cells were then collected by centrifugation at 3000 RCF for 15 min.

### Purification of the RRM-tandem constructs

Cell pellets were re-suspended in a lysis buffer containing 20 mM Tris−HCl, pH 7.5, 2 M NaCl, 25 mM imidazole, 0.2 mM TCEP, 1 mM PMSF, 0.5 mg/mL lysozyme, and 1 tablet of protease inhibitor. Cell lysate was sonicated after 3 freeze-thaw cycles and centrifuged at 23,710 RCF for 40 min using a Beckman Coulter Avanti JXN26/JA20 centrifuge. The supernatant was loaded onto 5 mL of HisPur Ni-NTA resin and washed with 200 mL of 20 mM Tris−HCl, pH 7.5, 2 M NaCl, 25 mM imidazole, and 0.2 mM TCEP. The sample was then eluted with 30 mL of 20 mM 2-Morpholinoethanesulfonic acid sodium salt (MES), pH 6.5, 500 mM imidazole, 500 mM Arg/Glu, and 0.2 mM TCEP. The eluted sample was cleaved with 2 μg/mL Ulp1 for 2 h at 25 °C (This step was skipped during the purification of SUMO tagged RBD and its RRM1 and RRM2 mutants) and diluted by threefold with a buffer A of 20 mM MES, pH 6.0, 100 mM Arg/Glu, and 0.1 mM TCEP before loading onto a 5-mL HiTrap Heparin column. The sample was eluted over a gradient with a buffer B of 20 mM MES, pH 6.0, 100 mM Arg/ Glu, 0.1 mM TCEP, 2 M NaCl, and 0.02% NaN3. RBD constructs were eluted around 50% B. Fractions containing the target proteins were pooled, concentrated, and loaded onto a HiLoad 16/600 Superdex 75 pg size exclusion column equilibrated with 20mM Tris-HCl pH 6.5, 0.4 M Arg/Glu, and 0.2mM TCEP.

### Purification of SRSF1

Cell pellets were re-suspended in a lysis buffer containing 20 mM Tris−HCl, pH 7.5, 150 mM Arg/Glu 2 M NaCl, 25 mM imidazole, 0.2 mM TCEP, 1 mM PMSF, 0.5 mg/mL lysozyme, and 1 tablet of protease inhibitor. Cell lysis followed by Ni-purification and SUMO cleavage were carried out in the same manner as the purification of RRM-tandem constructs. Then the protein eluted from the Ni column was diluted by threefold with a buffer A of 20 mM MES, pH 4.6, 100 mM Arg/Glu, and 0.1 mM TCEP before loading onto a 5-mL HiTrap Heparin column. The sample was eluted over a gradient with a buffer B of 20 mM MES, pH 11.5, 400 mM Arg/ Glu, 0.1 mM TCEP, 2 M NaCl, and 0.02% NaN3 in which the protein was eluted around 85% B. Fractions containing the target proteins were pooled, and pH was adjusted to 6.0-8.0 followed by concentration. The sample was exchanged to a buffer containing 20mM Tris-HCl pH 7.5, 0.80 M Arg/Glu, 0.2 mM TCEP and concentrated.

### Purification of RS tail constructs

Cell pellets were re-suspended in a lysis buffer containing 20 mM Tris-HCl, pH 7.5, 25 mM imidazole, 0.1 mM TCEP, and 6M Guanidinium HCl. Cell lysate was sonicated after 3 freeze-thaw cycles and centrifuged at 23,710 RCF for 40min using a Beckman Coulter Avanti JXN26/JA20 centrifuge. The supernatant was loaded onto 5 mL of HisPur Ni-NTA resin and washed with 200 mL of 20 mM Tris-HCl, pH 7.5, 0.2 mM TCEP, and 25 mM imidazole. The sample was then eluted with 30 mL of 500mM Arg/Glu, pH 7.0, 500 mM Imidazole, 1 mM TCEP, and 1 protease inhibitor cocktail. The eluted sample was cleaved with 2 μg/mL Ulp1 for 1 hour at 37 °C and diluted by twofold with a buffer A of 20mM sodium acetate, pH 5.0, 0.2 mM TCEP before loading onto a 5-mL HiTrap SP column. The protein was eluted over a gradient with a buffer B of 20 mM sodium acetate, pH 5.0, 2 M NaCl, and 0.2 mM TCEP. RS tail constructs were eluted approximately 30% B. Fractions containing the target proteins were pooled and concentrated. The concentrated proteins were exchanged and concentrated into a buffer containing 20 mM Tris-HCl, pH 7.5, and 0.2 mM TCEP.

### CD spectroscopy

CD spectroscopy was employed to test the conformation of folded RNA using a JASCO J-815 CD spectrometer. CD spectra of all folded 21-mer RNA (5 μM) were collected at 25 ^◦^C in 25 mM KCl, 0.25 M Arg/Glu, 20 mM Tris-HCl, pH 7.5, 0.1 mM EDTA, 0.1 mM TCEP, and 0.02 % TWEEN 20. Another set of CD spectra of the folded ARPC2 GQ RNA was collected at 25 ^◦^C and 95 ^◦^C in a 25 mM KCl without TWEEN 20. The CD spectrum of the 10 μM 8G RNA (unfolded) was collected in a 35 mM potassium phosphate buffer (35 mM K_3_PO_4_, 2.5 mM NaCl, 15 mM HEPES pH 7.5, 0.015% TWEEN 20, and 0.075 mM TCEP).

CD spectra were collected for the ARPC2 GQ RNA sample at a concentration of 5 μM, in the presence of varying concentrations of the RRM tandem (ranging from 2.5 to 15 μM). All the CD spectra were scanned three times to obtain spectral average using a 10-mm path length cuvette and a scanning speed of 50 nm/min, covering a wavelength range of 200–320 nm. Corresponding buffer spectra were subtracted for baseline correction.

### UV melting

UV melting and annealing experiments were conducted using 10-mm path length quartz cells on a JASCO V-730 spectrophotometer with a sample volume of 1.5 mL. The quartz cells were sealed to prevent the sample evaporation. Folded RNA was diluted to 5 μM in a buffer containing 25 mM KCl, 0.25 M Arg/Glu, 20 mM Tris-HCl, pH 7.5, 0.1 mM EDTA, and 0.1 mM TCEP. The absorbance at 295 nm was recorded after heating to 95 ^◦^C, followed by cooling to 25 ^◦^C at a 0.2 ^◦^C/min temperature gradient. The annealing and melting cycles were carried out in triplicate as three independent experiments.

### FP assays

Fluorescence assays for 8-mer RNA were collected in 35 mM K_3_PO_4_, 2.5 mM NaCl, 15 mM HEPES pH 7.5, 0.015% TWEEN 20, and 0.075 mM TCEP. Fluorescence polarization assays for 21-mer RNA were performed in 25 mM KCl, 0.25 M Arg/Glu, 20 mM Tris-HCl pH 7.5, 0.1 mM EDTA, 0.1 mM TCEP, and 0.02% TWEEN 20. Ten nM of 5’ fluorescein-labeled RNA was mixed with proteins at concentrations ranging from 8,000 nM to 0.488 nM, in black flat-bottom 96-well plates (Costar). Protein-RNA mixtures (50 μL) were shake at 100 RPM for 10 min, followed by incubation at 37 °C for 30 min, and then at 25 °C for 10 min. Fluorescence polarization measurements were carried out using a Cytation 5 with an excitation wavelength of 485 nm and an emission wavelength of 520 nm. We have shown that the total fluorescence intensities did not change during protein titration. Therefore, the dissociation constants (K_D_) were fitted using the quadratic equation with assumption of one-site interaction. The fluorescence polarization (Fp) binding curves were fitted using the quadratic equation below, where the fitting parameters F_min_, F_max_, and K_D_ represent the fluorescence polarization baseline, plateau, and dissociation constant, respectively. [P_T_] represents the total protein concentration, and [L_T_] represents the total RNA concentration (10 nM). Errors in dissociation constants were calculated based on three independent measurements.

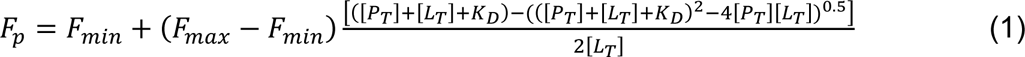

### Fluorescence spectroscopy

The Cy3-Cy5 labeled ARPC2 GQ RNA (200 nM) was dissolved in 100 μL of 25 mM KCl, 0.25 M Arg/Glu, 20 mM Tris-HCl, pH 7.5, 0.1 mM EDTA, and 0.1 mM TCEP. RS, the RRM tandem, KS, or RA stock solution (20 μM) dissolved in 20 mM Tris-HCl, pH 7.5, 0.4 M Arg/Glu, 0.1 mM TCEP was titrated into ARPC2 GQ RNA. Florescence emission spectra of protein-bound RNA were recorded from 530 nm to 700 nm using a Cytation 5 in black flat-bottom 96-well plates (Costar) at 25 °C. Full-length SRSF1 phase separates upon titrated into ARPC2 GQ RNA. To eliminate interference of phase separation, the SRSF1 RNA mixtures were centrifuged at 16,000 RCF for 10 min, and supernatants were used for fluorescence spectrum collection. Fluorescence emission spectra of the apo RNA were collected immediately after denaturation at 95 °C for 5 minutes. Samples were excited at 500 nm, and we have confirmed that this wavelength did not excite Cy5. All fluorescence spectra were collected in triplicate, and the average emission intensities were normalized by the Cy3 signal maximum at 566 nm after subtraction of baseline.

### Saturation Transfer Difference NMR

Mixtures of 500 μM 3-mer RNA oligomers and 10 μM RS were prepared in a D_2_O buffer with a pH of 5.6, supplemented with 1 mM MES. To eliminate residual H_2_O, the samples were freeze-dried and subsequently re-dissolved in 99.9% D_2_O. STD-NMR data were acquired at 298 K using an 850 MHz magnet and the sequence previously reported(56). A total of 16 transients were accumulated for all experiments.

### Plasmid design for dual luciferase assay, western blot and RT-PCR

The ARPC2 GQ DNA sequence was introduced upstream of the Renilla luciferase gene of the psiCHECK-2 plasmid via two separate mutagenesis polymerase chain reactions, with the first and second halves of the ARPC2 GQ introduced to the NheI restriction site downstream of the T7 promoter. The DNA encoding human SRSF1 was cloned into the pcDNA4 myc His A plasmid using BamH I and Hind III. Using mutagenesis polymerase chain reaction, pCDNA4 containing DNA encoding the RRM tandem was constructed.

### Cell culture and transfection

HEK 293 cells were cultured in Dulbecco’s Modified Eagles medium (DMEM) containing 10% fetal bovine serum (FBS) and 1% penicillin and streptomycin. Cells were grown at 37 °C in an incubator containing 5% CO_2_ and passaged into new media in a 6-well plate every 2-3 days. Cells were inoculated in 96 well-plates prior to the transfection for the dual Luciferase assay. Once the cells reached to 60–70% confluency, cells were transfected with wild-type psiCHECK-2, the psiCHECK-ARPC2, pCDNA4-FL SRSF1, or pCDNA4-RBD using LipofectAMINE 3000 (Invitrogen) according to the manufacturer’s instructions. After transient transfection, cells were incubated at 37°C for 48 h until the cell density reached a confluency of about 95%.

### Dual luciferase assay

Dual luciferase assays were conducted after lysing the transfected cells using the Dual-luciferase Reporter Assay kit (Promega) according to the manufacturer’s instructions. The firefly and Renilla luciferase activities of each tested condition were measured in three independent replicates using a Cytation 5 plate reader. The average luciferase activity was determined as the ratio of Renilla to firefly luciferase activity. The Renilla/firefly luminescence ratio of psiCHECK2 was used to normalize other datasets.

### Western Blot

Western blots were carried out after the lysis of the transfected cells with SDS-PAGE loading buffer. Western blots were conducted as previously described(57). SRSF1 and the RRM tandem were immunoblotted with a primary antibody called Goat anti-ECS (DYKDDDDK) and Anti Goat, a secondary antibody conjugated with horseradish peroxidase (HRP). Antigen–antibody complexes were visualized by the SYNGENE gel imager and GeneSys gel imaging software.

## Results

### The RS domain of SRSF1 demonstrates RNA binding specificity

Given that SRSF1 RS contains 19 arginine residues, it is intuitive to conceive that the RS domain can contact nucleotides via electrostatic interactions. However, the RNA-binding specificity of RS has never been examined. To this end, we used fluorescence polarization (FP) to measure the binding affinities of RS to various RNA oligonucleotides (Fig. 1A). To prevent protein RNA co-aggregation, which may interfere with binding assays, we carefully chose the RNA lengths and salt concentrations.

We found that 8-mer RNA did not aggregate in a buffer containing 35 mM potassium phosphate (Fig. S1A). In searching for the optimal salt concentration for binding assays, we found that salts significantly weakened the interaction between RS and RNA, suggesting a role of electrostatic interactions in RS and RNA binding (Fig. S1B). Of all tested RNA, RS had the highest affinity to 8-mer poly-G. We found that 8-mer poly-G formed intermolecular G-quadruplex, as shown by the diagnostic negative peak at 240 nm and the positive peak at 267 nm (Fig. S1C). Therefore, its binding affinity to RS cannot be compared with other 8-mer single-stranded RNA due to differences in structure and net charges. As a substitute for 8-mer poly-G, we tested 8-mer GA, which is unable form GQ. Apart from 8-mer poly-G, RS bound more tightly to 8-mer GA than to 8-mer CU or other polymers (Fig. 1B, Table 1). Therefore, we concluded that RS preferred purine rich sequences. As the nucleotides tested here had the same number of backbone phosphates, the difference in RS binding affinity was attributed to nucleobases.

**Table 1:** RS prefers purine-rich RNA.

| Nucleotide | K<sub>D</sub> (nM) |
| --- | --- |
| 8G* | 3.0 ± 0.8 |
| 8A | 613 ± 15 |
| 8C | 1220 ± 40 |
| 8U | 550 ± 21 |
| 4CU | 470 ± 12 |
| 4GA | 83 ± 3 |

\* 8G RNA forms an intermolecular G-quadruplex, making FP values higher than other RNA oligomers in the single-stranded conformation.

### Both the RS and the tandem RRM are essential for RNA GQ binding of SRSF1

The 5’ untranslated region of human ARPC2 mRNA contains a 21-nucleotide sequence that forms G-quadruplex (ARPC2 GQ) (Fig. 2A). A previous study has found that ARPC2 GQ can pull down SRSF1, along with other SR proteins (55). Our success in obtaining soluble SRSF1 gave us an opportunity to quantitatively examine this interaction and to investigate which SRSF1 domains are responsible for the GQ binding. To this end, we carried out FP assays in a buffer that contained 25 mM KCl and 250 mM Arg/Glu. Arg/Glu was used to prevent phase separation, which may interfere with FP assays or circular dichroism (CD) characterization. We confirmed that the ARPC2 GQ RNA was successfully folded into G-quadruplex in the presence of 25 mM KCl, as suggested by the negative peak around 240 nm and the positive peak around 267 nm in the CD spectrum (Fig. 2B). The presence of Arg/Glu did not change the CD spectrum of the ARPC2 GQ (Fig. S2A). The formation of GQ was further confirmed by the characteristic UV melting curve monitored at 295 nm (Fig. S2B).

**Figure 2:**
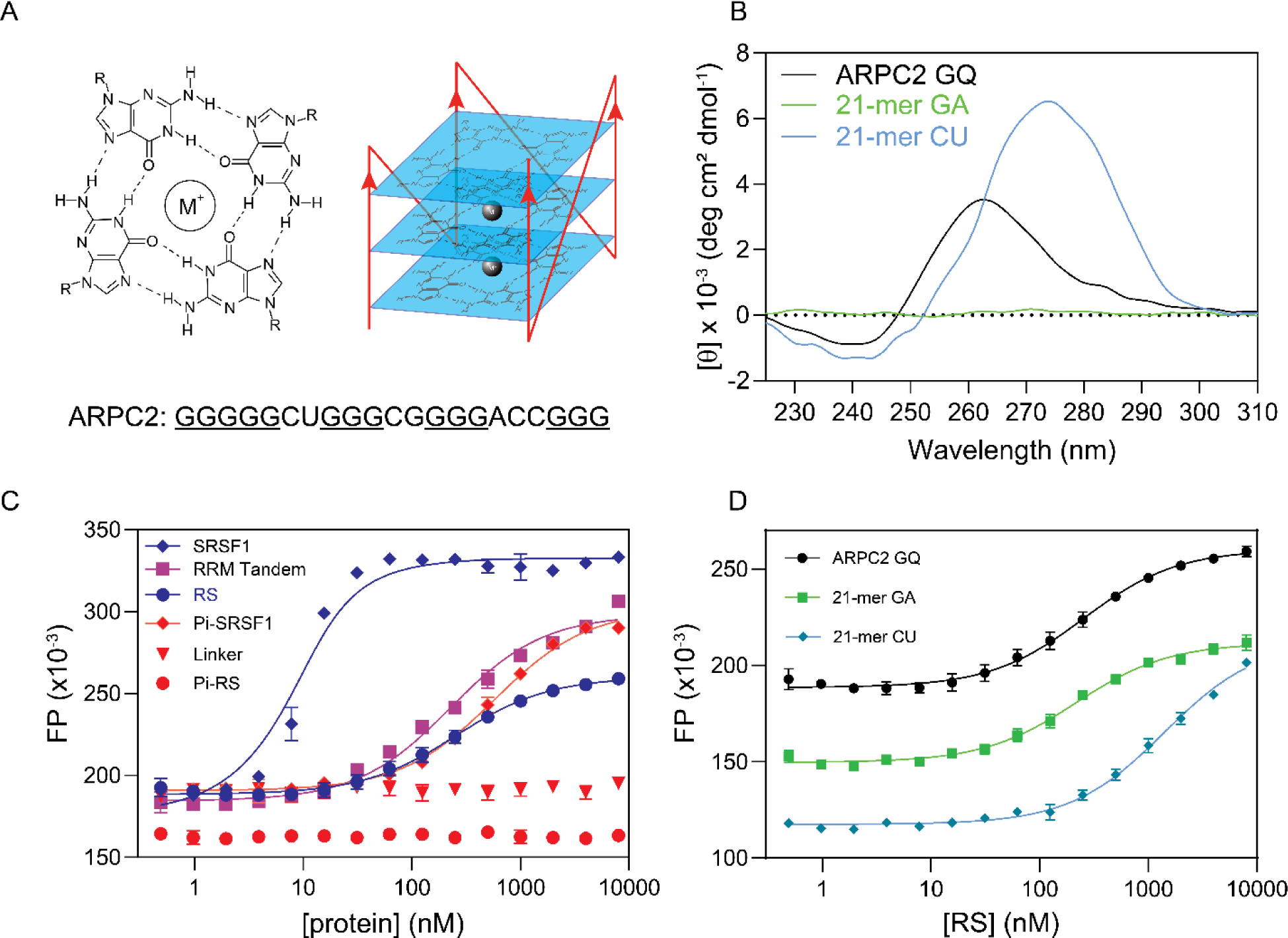
Both RS and the RRM tandem of SRSF1 are essential for its interaction with the ARPC2 GQ. (A) Schematic diagram of G-quadruplex. M^+^ denotes mono-ions. A parallel GQ is shown on the right. Mono-ions lay between G-tetrad layers and form bonds with carbonyl oxygen atoms. The ARPC2 GQ sequence used in this study is shown below. Putative guanine residues that are involved in forming G-tetrads are underlined. (B) The circular dichroism spectrum of the ARPC2 GQ. (C) FP binding curves of the ARPC2 GQ with various SRSF1 constructs. (D) FP binding curves for RS with 21-mer GA, 21-mer CU, and the ARPC2 GQ. FP measurements were performed in a buffer containing 25 mM KCl, 0.25 M Arg/Glu, 20 mM Tris-HCl pH 7.5, 0.1 mM EDTA, 0.1 mM TCEP, 0.02 % TWEEN 20. Fluorescein was attached to the 5’ end of RNA, and the probe concentration was 10 nM for all samples.

In this buffer, SRSF1 bound with ARPC2 GQ with a K_D_ of 4 nM (Fig. 2C, Table 2). RS or RBD alone bound to the GQ RNA with a much lower affinity (K_D_ is around 200 nM). Whereas, the RRM1/RRM2 linker has no detectable binding to the ARPC2 GQ (Fig. 2C). These results together suggested that both the RRM tandem and RS were necessary for the ARPC2 GQ binding. As RS is subject to phosphorylation in the cell, we further tested how phosphorylation affected its GQ binding and found that hyperphosphorylation of RS completely abolished its binding to GQ. Different from isolated RS, hyper-phosphorylated SRSF1 binds to ARPC2 GQ with a lower affinity than the RRM tandem (K_D_ of 570 nM versus 220 nM), suggesting that phosphorylation not only abolishes RS RNA binding, but also inhibits RNA binding of the neighboring RRM tandem.

**Table 2:** Both the RRM tandem and the RS domain of SRSF1 are essential for its interaction with the ARPC2 GQ.

| Protein constructs | RNA | K<sub>D</sub> (nM) |
| --- | --- | --- |
| SRSF1 | ARPC2 GQ | 3.6 ± 0.7 |
| RS | ARPC2 GQ | 250 ± 22 |
| RRM Tandem | ARPC2 GQ | 220 ± 19 |
| RRM1/RRM2 linker | ARPC2 GQ | No binding |
| Pi-SRSF1 | ARPC2 GQ | 570 ± 35 |
| Pi-RS | ARPC2 GQ | No binding |
| RS | 21-mer GA | 210 ± 16 |
| RS | 21-mer CU | 1400 ± 100 |

**Table 3:** The RRM tandem uses its classic RNA-binding sites for its interaction with the ARPC2 GQ.

| RRM tandem mutants | K<sub>D</sub> (nM) | Relative K<sub>D</sub> |
| --- | --- | --- |
| WT | 130 ± 11 | 1 |
| RRM1 Y19A | 1,800 ± 160 | 14 |
| RRM1 F56A | 7,000 ± 1,500 | 54 |
| RRM1 F58A | 1,500 ± 80 | 12 |
| RRM2 W134A | 1,200 ± 90 | 9 |
| RRM2 Q135A | 350 ± 20 | 3 |
| RRM2 R173A | 140 ± 12 | 1 |

Using 8-mer RNA, we showed that RS preferred purine-rich RNA. We wondered whether this RNA-binding preference remained for longer RNA. To this end, we compared binding affinities to ARPC2 GQ (21-mer), poly-GA, or CU of the same length. Consistent with our findings using 8-mer RNA, the binding affinity of RS to 21-mer GA is 7-fold higher than that to 21-mer CU (K_D_ of 210 +/-16 nM versus 1400 +/-100 nM). RS binds to the ARPC2 GQ with a similar K_D_ to 21-mer GA. Notably, 21-mer GA did not form GQ, according to our CD data (Fig. 2D). These results suggest that RS prefers guanine rich RNA sequences regardless of their secondary structure.

### Identification of key SRSF1 residues responsible for ARPC2 GQ binding

We have shown that both the RRM tandem and RS are required for ARPC2 GQ binding. To understand the binding mechanism, we wanted to identify the SRSF1 residues essential for ARPC2 GQ binding. RRM1 and RRM2 of SRSF1 recognize C- and GGA-containing single-stranded RNA motifs, respectively (25,26). These motifs are also present in ARPC2 GQ. Others and our studies have identified the SRSF1 residues for RNA binding, such as Y19, F56, F58, W134, and Q135 (Fig. 3A) (26,28). In our previous study, we have confirmed that mutation of these residues does not perturb the protein structure (28). Therefore, we compared binding affinities of the wild-type RRM tandem with its mutants. R173 of RRM2 is distal to its RNA-binding site and was selected as a negative control. We found that the SRSF1 residues essential for binding to classic RNA ligands were also important for ARPC2 GQ binding (Fig. 3B), suggesting that the RRM tandem recognizes ARPC2 GQ via the binding sites for single-stranded RNA (ss-RNA). These residues form either stacking interactions (Y19, F56, F58, W134) or H-bonds (Q135) with nucleobases of ss-RNA.

**Figure 3:**
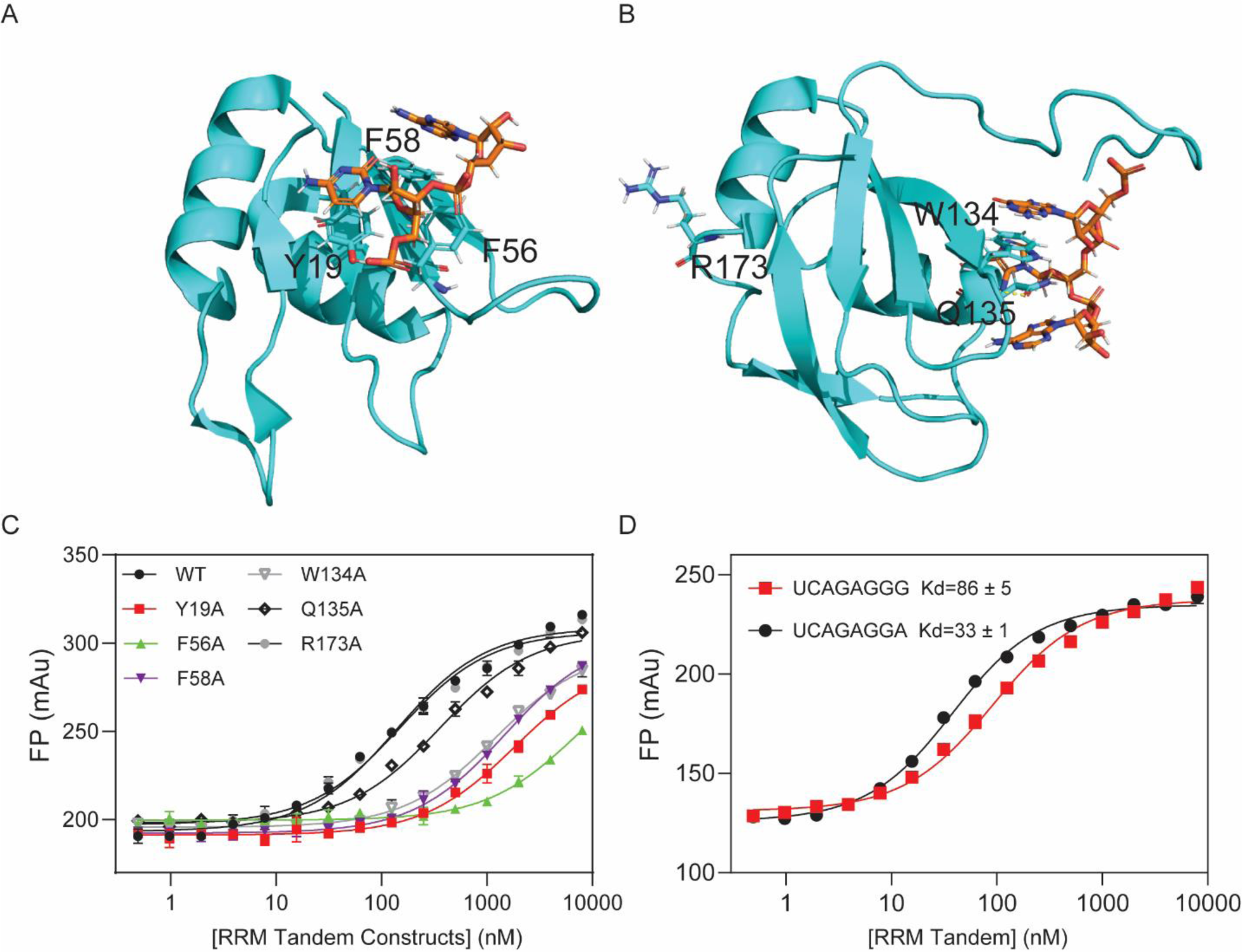
RRM1 and RRM2 residues for recognizing single-stranded RNA are also essential for ARPC2 GQ binding. (A) Structure of RRM1 (PDB ID: 6HPJ) and (B) RRM2 (PDB ID: 2M8D) with their cognate RNA motifs. RNA and protein residues are shown by orange and cyan sticks, respectively. Molecular graphics were prepared using PyMOL. (C) FP binding curves of ARPC2 GQ with the RRM tandem and its mutants. (D) FP binding curves of the RRM tandem with GGA and GGG motifs.

GQ-forming sequences usually contain at least 4 G-tracts of 3 consecutive guanines. Therefore, it is of interest to test whether the SRSF1 tandem RRMs binds with the GGG motifs, in addition to its optimal motif GGA. Therefore, we used the GGG motif to replace the GGA motif of UCAGAGGA, one of classic consensus RNA ligands for SRSF1. We found that replacement of GGA by GGG only reduced binding affinity by three-fold. Therefore, it is possible that RRM2 either binds to GGA or GGG if these sites are exposed. In summary, our results suggest that the SRSF1 RRM tandem uses its classic RNA-binding sites to bind with C- and GGA-/GGG-containing motifs.

In the previous sections, we showed that RS demonstrated RNA-binding specificity, and this specificity is unlikely attributed to charge-charge interaction. SRSF1 RS contains 19 arginine residues and 20 serine residues, accounting for three quarters of its amino acid composition. We wondered what residues are responsible for the RS and ARPC2 GQ binding. To this end, we created two RS mutants, KS and RA, in which all arginine or serine residues were replaced by lysine (the KS mutant) or alanine (the RA mutant), respectively (Fig. 4A). The KS mutant has the same charge distribution as RS. However, its binding to ARPC2 GQ was too weak to be detected (Fig. 4B). In contrast, RA showed a similar RNA-binding affinity to RS (Fig. 4B). These results suggest that arginine residues in RS are the main contributors to its GQ binding.

**Figure 4:**
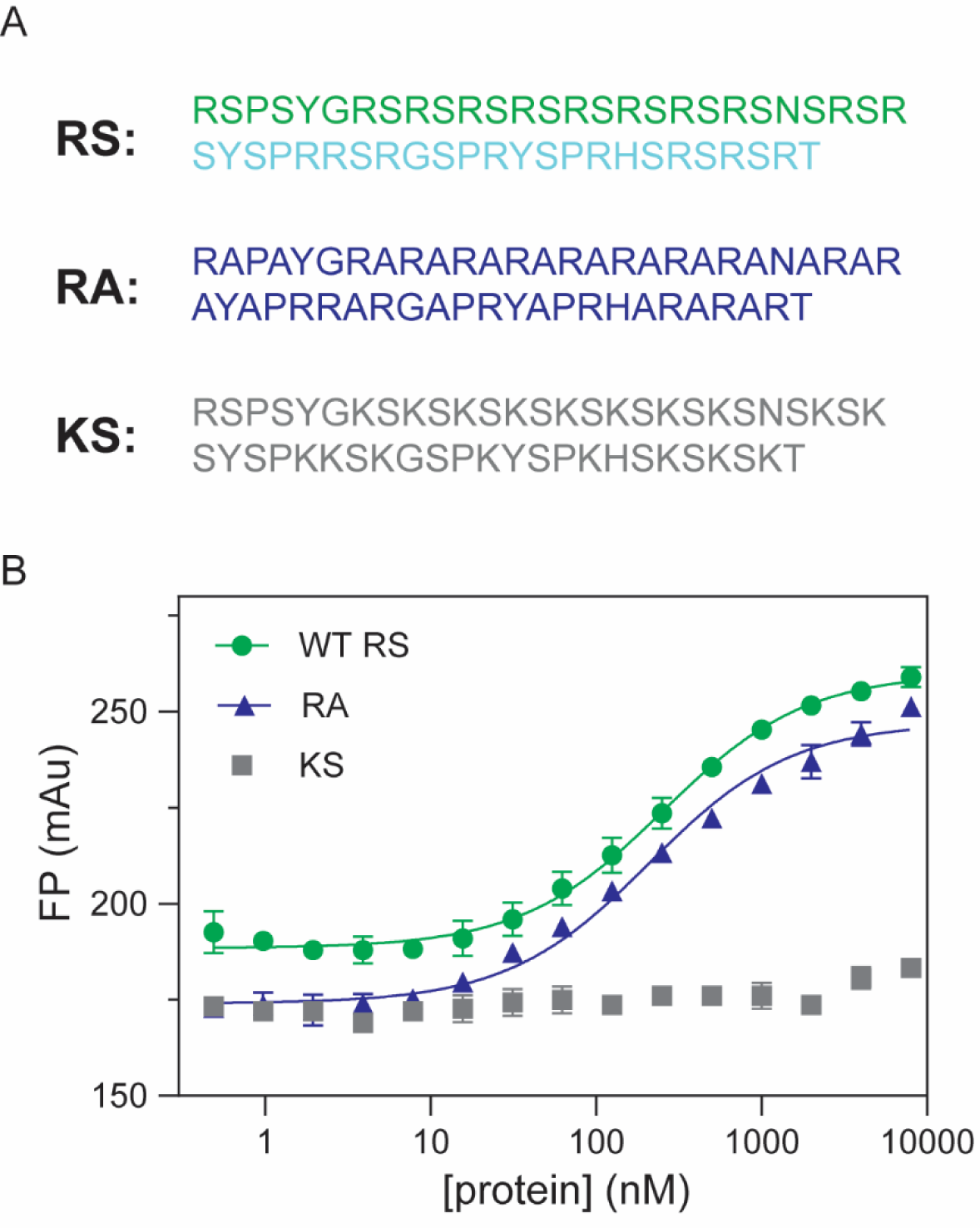
Arginine residues in RS are the main contributors for ARPC2 GQ binding. (A) Sequences of RS and its mutants, RA, and KS. ARPC2 GQ(10 nM) is labeled by fluorescein at the 5’ end. (B) FP binding curves for ARPC2 GQ and RS, KS, and RA.

### SRSF1 unfolds ARPC2 GQ

RNA GQ is folded in such a way that the majority of guanine nucleobases are packed inside, while the phosphate backbones are exposed (Fig. 2A). However, the RNA-binding specificity of the RRM tandem and RS requires nucleobases to be exposed. This implies that SRSF1 binding potentially unfolds the GQ. To test this hypothesis, we attached the 5’ and 3’ ends of ARPC2 GQ RNA with Cy5 and Cy3 fluorescent probes, respectively (Fig. 5A). The formation of GQ allows fluorescence resonance energy transfer (FRET) between Cy3 and Cy5, which have a Förster range of 53 Å (58). We observed the Cy5 fluorescence emission at 670 nm when the sample was excited with light at 500 nm. We also confirmed that Cy5 alone was not excited by light at 500 nm (Fig. 5A). Therefore, the observed FRET is attributed to GQ formation. To confirm that the Cy5 signal can be used to probe unfolding of GQ, we also collected a spectrum for the thermally denatured GQ, which has a lower Cy5 signal (Fig. 5A). To rule out the possibility that Cy5 or Cy3 labeling affects GQ formation, we collected CD spectra and confirmed that fluorophore labeling did not disturb the GQ structure (Fig. S3A).

**Figure 5:**
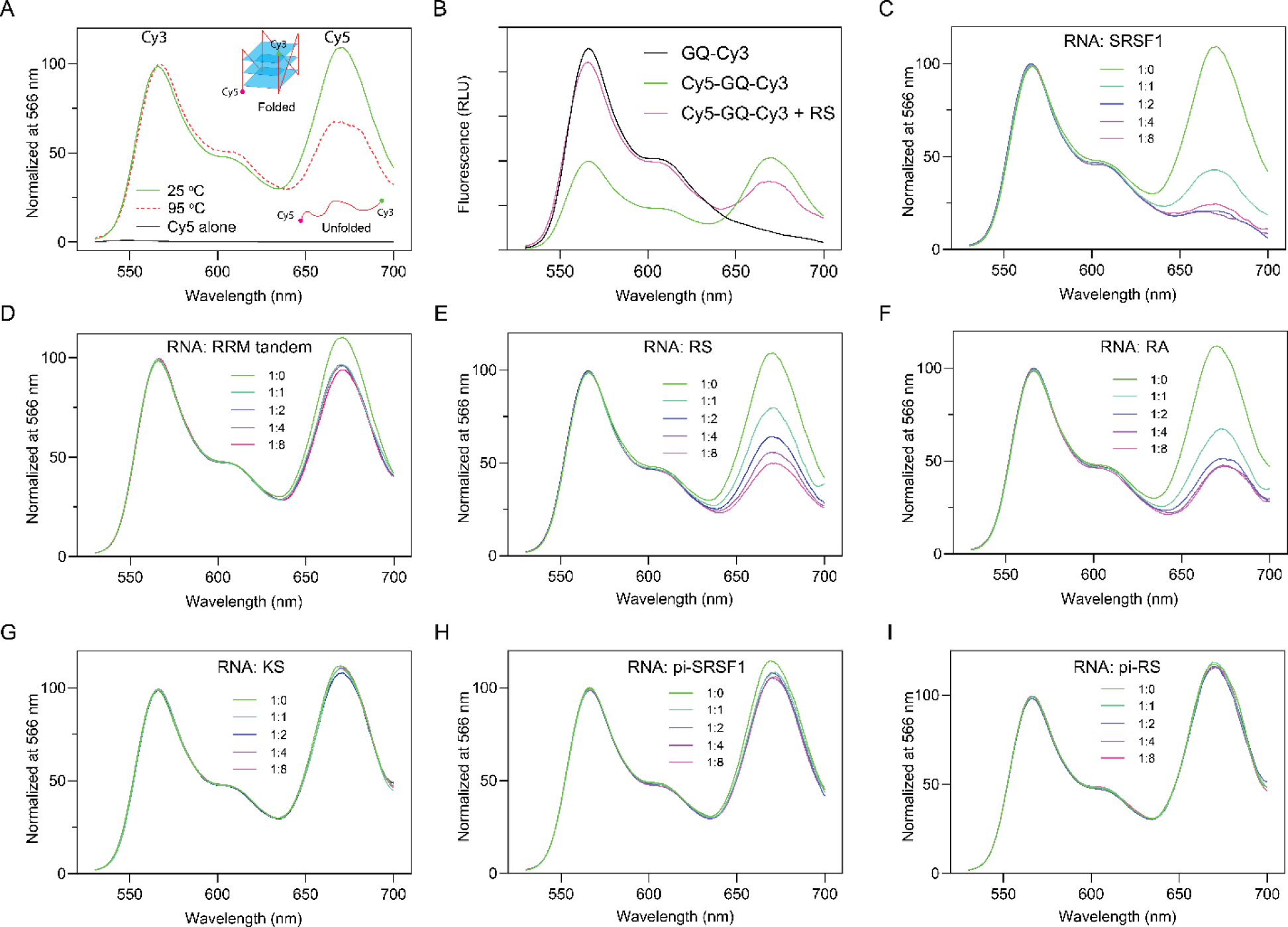
The RS domain of SRSF1 plays a dominant role in unfolding ARPC2 GQ. (A) The principle of the experimental design. Cy5 and Cy3 were labeled at the 5’ and 3’ ends of ARPC2 GQ, respectively. When excited by light at 500 nm, the fluorescence spectrum of ARPC2 GQ demonstrated the Cy3 signal maxima at 566 nm and the Cy5 signal maxima at 670 nm due to FRET (green line). Thermally denatured GQ has a relatively lower Cy5 signal. Cy5 alone could not be excited by light at 500 nm (black line). (B) Control experiments to test whether RS quenches Cy3. The fluorescence spectra in the relative light unit (RLU) were normalized to the same RNA concentrations. Fluorescence spectra were collected when ARPC2 GQ was titrated with (C) full-length SRSF1, (D) the RRM tandem, (E) RS, (F) RA, (G) KS, (H) phosphorylated SRSF1 (pi-SRSF1), and (I) phosphorylated RS (pi-RS). All spectra were collected with excitation light at 500 nm and the ARPC2 GQ concentration is 200 nM.

To avoid interference from potential phase separation or aggregation, we dissolved ARPC2 GQ and SRSF1 in a high ionic strength buffer containing 25 mM KCl and 0.25 M Arg/Glu and centrifuged the samples before collecting fluorescence data. This experiment design depends on the relative fluorescence intensity decrease of Cy5 to detect GQ unfolding. To rule out the possibility of fluorescence quenching by RS, we performed two control experiments. In the first control experiment, we compared the fluorescence spectra of Cy3- and Cy3/Cy5-labeled ARPC2 GQ. The Cy3 intensity of the single-labeled ARPC2 GQ is higher than that of double-labeled RNA (Fig. 5B, blank versus green lines), as the Cy3 energy is transferred to Cy5 via FRET in the double-labeled RNA. Upon addition of RS (Fig. 5B, purple line), the Cy5 signal decreases and the Cy3 signal increases relative to double-labeled the RNA GQ. These results suggest there is no signal quenching for Cy3. We further compared fluorescence intensity of Cy5 with and without RS and found that addition of RS enhanced instead of quenching the Cy5 signal (Fig. S3B).

An obvious decrease in the Cy5 signal was observed with gradual titration of SRSF1 (Fig. C). It is noteworthy that the Cy5/Cy3 ratio for SRSF1-bound ARPC2 GQ was lower than that of the sample denatured at 95 °C (Fig. 5A), suggesting that SRSF1 is more efficient in unfolding ARPC2 GQ than thermal denaturation. We further investigated the roles of the RRM tandem and RS in unfolding GQ. We found that the RRM tandem decreased the Cy5 relative signal by less than 20% (Fig. 5D). This is consistent with our circular dichroism results, which showed a mild decrease in the GQ-characteristic peak around 267 nm (Fig. S3C). In contrast, RS unfolds ARPC2 GQ more efficiently (Fig. 5E). In line with our binding assays (Fig. 4B), the RA mutant can unfold ARPC2 GQ (Fig. 5F), whereas KS cannot. (Fig. 5G). As a negative control, we also tested BSA and found that it was unable to unfold ARPC2 GQ (Fig. S3D). Furthermore, phosphorylated SRSF1 (Fig. 5H) or RS (Fig. 5I) has lower efficiency in unfolding ARPC2 GQ, which agrees with our FP binding assays (Fig. 2A).

### The Arg sidechain interacts with the Guanidine base, which may be responsible for GQ unfolding

In previous sections, we have shown that RS has RNA-binding specificity, and Arg residues in RS play a main role in unfolding ARPC2 GQ. In addition to forming electrostatic interactions with phosphate backbones, arginine residues are also able to form H-bonding and stacking interactions with nucleobases. As RS unfolds GQ, the complex may exist in numerous conformations, which makes crystallography unsuitable. NMR is a powerful method to characterize dynamic interactions. However, RS and ARPC2 GQ phase separate at the concentration for NMR studies (> 50 μM), making our system a difficult case. To avoid phase separation, we tested RNA homopolymers of different lengths and found that 3-mer polynucleotides remained soluble after being mixing with RS. This success enabled us to use saturation transfer difference (STD) NMR to characterize the interaction between RS and homo-polymeric RNA. STD NMR selectively saturates resonances of a functional group of one binder (usually large molecules). The saturation can be diffused to neighboring protons through intramolecular proton-proton cross relaxation. Due to transient interactions, the saturation can be transferred to the ligand via inter-molecular NOE, which enables the identification of epitopes involved in binding (Fig. 7A). STD is a sensitive method to detect weak binding with a K_D_ up to the millimolar range. If Arg sidechains interact with nucleobases, STD signal should be observed when saturation is applied to Arg sidechains. We irradiated methylene groups of Arg sidechains, whose NMR chemical shifts are clustered around 2.5 ppm, away from the chemical shift of nucleobases (> 7.0 ppm). We collected STD data for 3-mer poly-A, C, G, and U with sub-stoichiometric amount of RS. As shown by Fig. 6B, only GGG of the four 3-mer RNAs showed noticeable STD signals. These results suggested that the guanine base was involved in interacting with the Arg sidechain but others. Our results also agree with a previous bioinformatic study that shows that the interaction between the Arg sidechain and guanine ranks the most probable one among protein DNA recognition(59).

**Figure 6:**
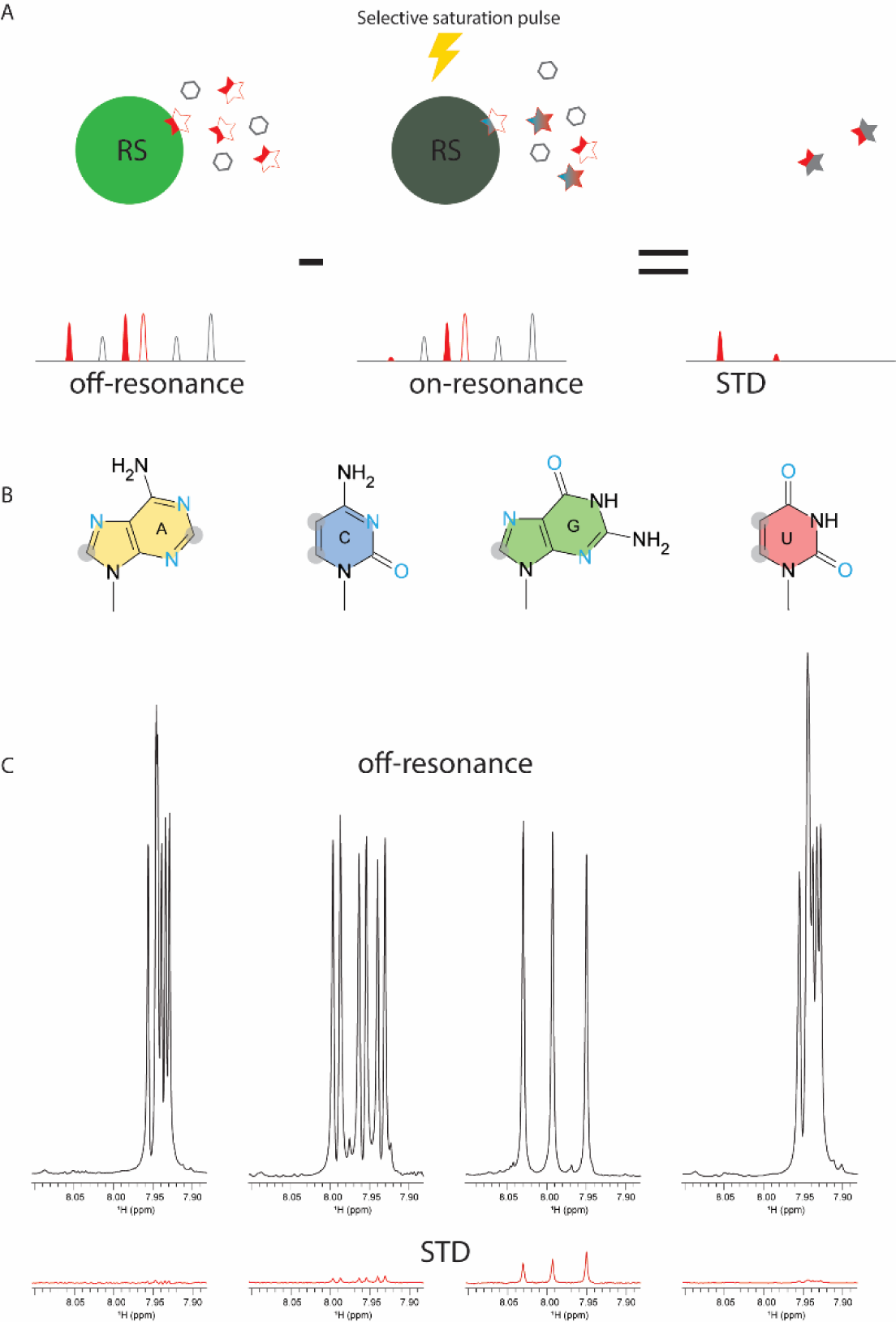
Arg sidechain interacts with guanine. (A) Principle of saturation transfer difference (STD) NMR. NMR spectra are collected with and without the saturation pulse focused on the protein region that is away from the ligand resonance. Due to binding and exchange, the NMR signal of the protons directly involved in binding or adjacent to binding sites will be saturated. Subtraction of the saturated spectrum from the un-saturated one yields the STD spectrum, in which only the protons adjacent binding sites survive, and the rest canceled out. (B) Structure of nucleobases, with nonexchangeable C-H groups shadowed by gray circle and H-bonding acceptors in blue. (C) STD NMR of 3-mer RNA (0.5 mM) with sub-stoichiometry amount of RS (10 uM). The reference NMR spectra without saturation were shown in black and the STD spectra shown in red. The samples were dissolved in D2O solution to avoid saturation transfer through solvent. (C) The schematic illustration of interactions between the Arg guanidino group and guanine. Note that guanine the only nucleobase that has two sites serving as H-bonding acceptors.

**Figure 7.**
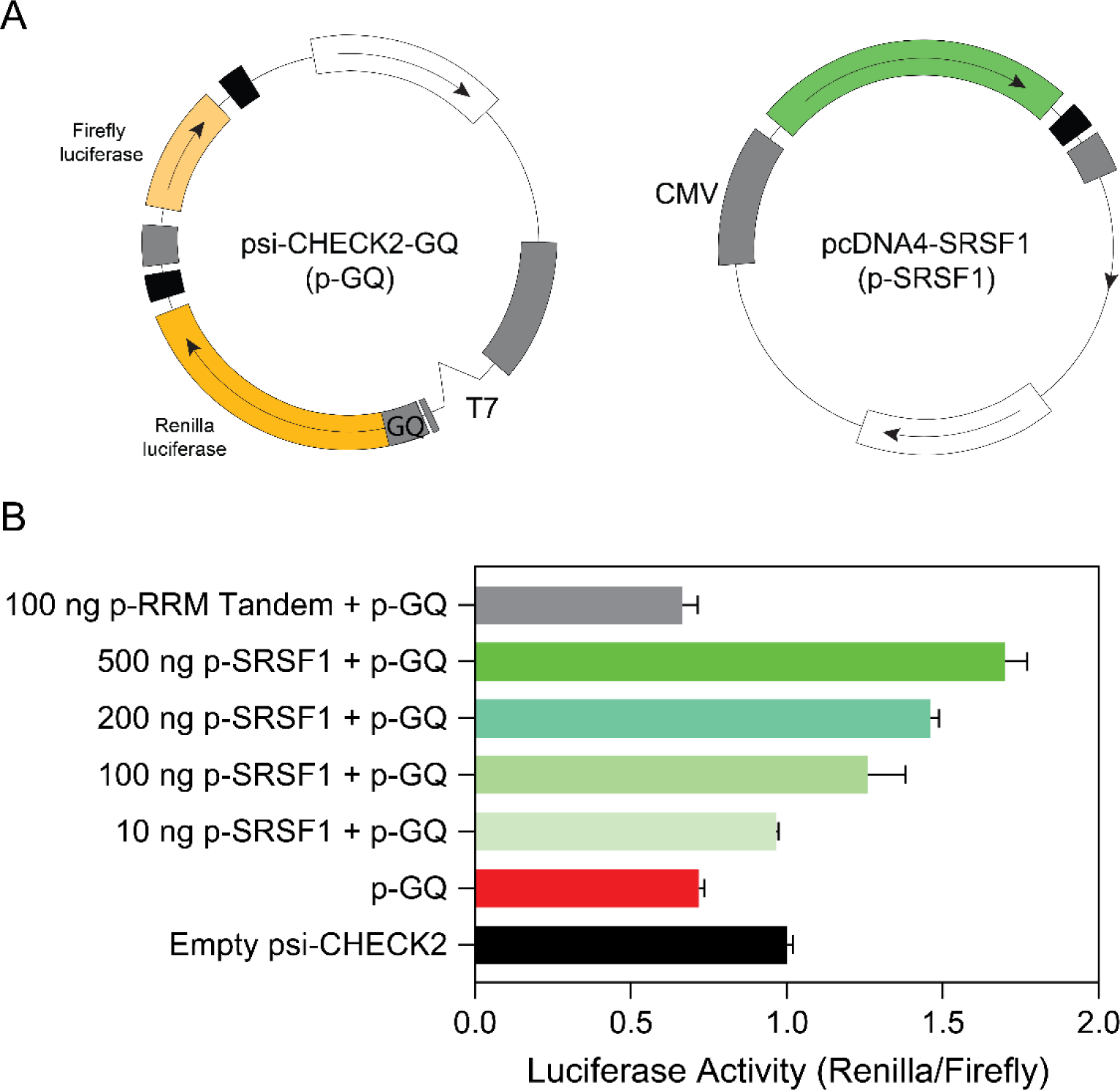
SRSF1 is able to rescue the gene expression inhibited by the ARPC2 GQ. (A) Plasmid design of psi-CHECK2-GQ (GQ). The ARPC2 GQ RNA sequence (21 nucleotides) was inserted immediately before the starting codon of the Renilla luciferase gene and after the T7 promoter. The empty psi-CHECK2 vector contains a G-rich sequence that is unable to form GQ. (B) Renilla and firefly luciferase activity ratio for HEK293 cells transfected by empty psi-CHECK2 (100 ng), or psi-CHECK2-GQ (100 ng), or co-transfected by psi-CHECK2-GQ and pcDNA4 plasmid harboring SRSF1 constructs. All assays were performed three times for error estimation. The luciferase luminescence was recorded by a Cytation 5 plate reader.

### SRSF1 alleviates translation inhibition induced by the ARPC2 GQ

The presence of GQs in mRNA typically impedes gene expression by creating steric hindrance. Previous studies have used dual luciferase assays to detect a 40% reduction in translation efficiency in the presence of ARPC2 RNA GQ(55). We hypothesized that if SRSF1 can unwind RNA GQs within the cell, it should be capable of rescuing the reduction in gene expression caused by GQ formation. To investigate this hypothesis, we introduced the ARPC2 GQ sequence upstream of the Renilla luciferase gene in the psi-CHECK2 plasmid (Fig. 7A). The firefly luciferase gene within this plasmid served as a reference gene to standardize transfection efficiency. Additionally, we cloned SRSF1 or the RRM tandem into the pcDNA4 plasmid for co-transfection (Fig. 7A).

Upon transfection of HEK293 cells with the psi-CHECK2 plasmid containing the ARPC2 GQ sequence, the ratio of luciferase activity between Renilla and firefly was reduced to 70% compared to the psi-CHECK2 plasmid lacking the GQ-forming region (Fig. 7B). However, co-transfection of HEK293 cells with 10 ng of pcDNA4 encoding SRSF1 restored 96% of the luciferase activity. Notably, this rescuing effect displayed dose-responsive behavior (as depicted in Fig. 7B).

Importantly, the RS domain was identified as the key contributor to this rescue effect, as the RRM tandem alone failed to restore Renilla luciferase activity (Fig. 7B). We confirmed that the full-length SRSF1 and the RRM tandem were expressed using western blots (Fig. S4). It’s worth mentioning that the RS domain is inherently unstructured and susceptible to degradation and modification within the cell, which is why we did not evaluate the co-transfection of the RS-encoding plasmid in the luciferase assays.

## Discussion

RS domains are abundant in the human proteome, as exemplified by SR proteins and SR-like proteins. In SR proteins, the primary function of RS domains has been found to mediate protein-protein interactions. Due to their extensive electropositive charges, RS domains are generally considered to have nonspecific and, in some cases, undesirable roles as RNA binders, often regarded as secondary in most scenarios. Indeed, the RS domain of SRSF1 is unnecessary for splicing of RNA transcripts with robust splicing enhancers (26,60). Nevertheless, several studies have indicated that RS domains can play a role in RNA binding. Shen et al. have shown the crosslinking between RS domains and RNA transcripts (42,43). Moreover, RS domains are indispensable for RNA transcripts with weak splicing enhancers, such as IgM (42,46). Collectively, these studies suggest that RS enhances RNA binding of SRSF1. Our research has provided a new insight, revealing that RS domains exhibit binding preference for purine-rich RNA. Therefore, it is conceivable that RS domains not only enhance RNA binding but also fine tune the RNA-binding specificity of SRSF1.

The preference of RS for purine-rich RNA underpins its role in RNA GQ binding. A previous study has found that the ARPC2 GQ-forming region can pull-down most of SR family members (61). For SRSF1, the RRM tandem and the RS domain have an overlapping binding specificity, which is coincidental. RRM domains from other SR proteins have distinct RNA-binding specificity. For example, the SRSF3 RRM prefers pyrimidine-rich RNA (29,62-64). Therefore, it is unlikely that all SR proteins depend on their RRM domains to binding RNA GQ. On the contrary, RS domains of SR proteins are shared across the family, suggesting RS domains are probably responsible for GQ binding for some SR proteins. This is in line with the aforementioned findings of RS/RNA crosslinking and RS-dependent splicing for some RNA transcripts (42,43,46). In addition, we showed that phosphorylation completely abolishes RNA binding of RS. Therefore, others and our results together suggest a portion of SR proteins exist unphosphorylated in the cell so that RS domains can play their roles in RNA binding.

SRSF1 has the shortest RS among the 12 SR proteins. It is likely that RS domains play more important roles in GQ binding for other SR proteins. Among tens of thousands predicted GQ-forming RNA sequences, only a small fraction of them takes GQ conformation in the cell (52). In addition to RNA helicases that unwind RNA GQs, it is inferred that various RNA-binding proteins are responsible for preventing GQ formation in the cell (52). However, only a few proteins have been identified to have this function, such as hnRNPH1 (54). However, the scarcity of proteins binding with G-rich RNA cannot explain how GQ formation is prevented in the cell. Given the abundance of RS-containing proteins, it is likely that SR or SR-like proteins may be one of the main players that prevent GQ formation. Although SRSF1 RRM domains have a secondary role in unfolding GQ, they are important for tight binding to GQ. It is likely that SRSF1 RRMs help to locate the protein to purine-rich regions for the RS domain to unfold the GQ. The RRM domains from other SR proteins may locate the proteins to their individual cognate RNA motifs and their RS domains unfold neighboring GQ sequences.

We have discovered that interactions between Arg residues in RS domains and the guanine nucleobases play a pivotal role in binding (Fig. 4) and unfolding (Fig. 5) of GQs. The GQ structure is stabilized by H-bonds, base stacking, and chelation bonds with ions. To unfold GQ, a new set of interactions should be established to replace the interactions stabilizing GQ. The uniqueness of the Arg sidechain lies in its folk-like sidechain, which can form various interactions with nucleobases, including H-bonds, salt-bridges, and stacking interactions (Fig. 8A). Among the four nucleobases, guanine stands out due to its possession of three H-bonding acceptor sites that can form H-bonds with Arg sidechain in multiple ways (Fig. 8A). Notably, the Arg sidechain can simultaneously form hydrogen bonds with guanine while creating salt bridges with the phosphate backbone, as visualized in Fig. 8B. This dual interaction capability makes Arg/guanine interactions the most prevalent in DNA-binding and RNA-binding proteins according to two comprehensive bioinformatic analysis (59,65). Our STD NMR results have experimentally confirmed that the Arg sidechain specifically interacts with guanine, with no similar interactions observed for other nucleobases. These findings provide a mechanistic foundation for understanding the interactions between RS domains and G-rich sequences. According to our ensemble FRET results, RS-bound ARPC2 GQ takes a more extended conformation than the apo GQ unfolded by thermal denaturation. As thermal denatured polymers assume a random conformation, conformers that have Cy3 and Cy5 close enough for FRET still exist. However, RS bound ARPC2 GQ has a much lower FRET. This suggests a cooperative binding of SRSF1 RRMs and RS to ARPC2 GQ, which helps maintain the RNA in an extended conformation. It is noteworthy that in this extended conformation, the distance between neighboring backbone phosphates of the RNA ranges from 6.5 to 7.5 Å, closely matching the pitch of the RS dipeptide in the extended conformation (6.8 Å). Based on our experimental findings, we propose the model depicted in Fig. 8C to elucidate the mechanism by which RS domains unwind GQs.

**Figure 8:**
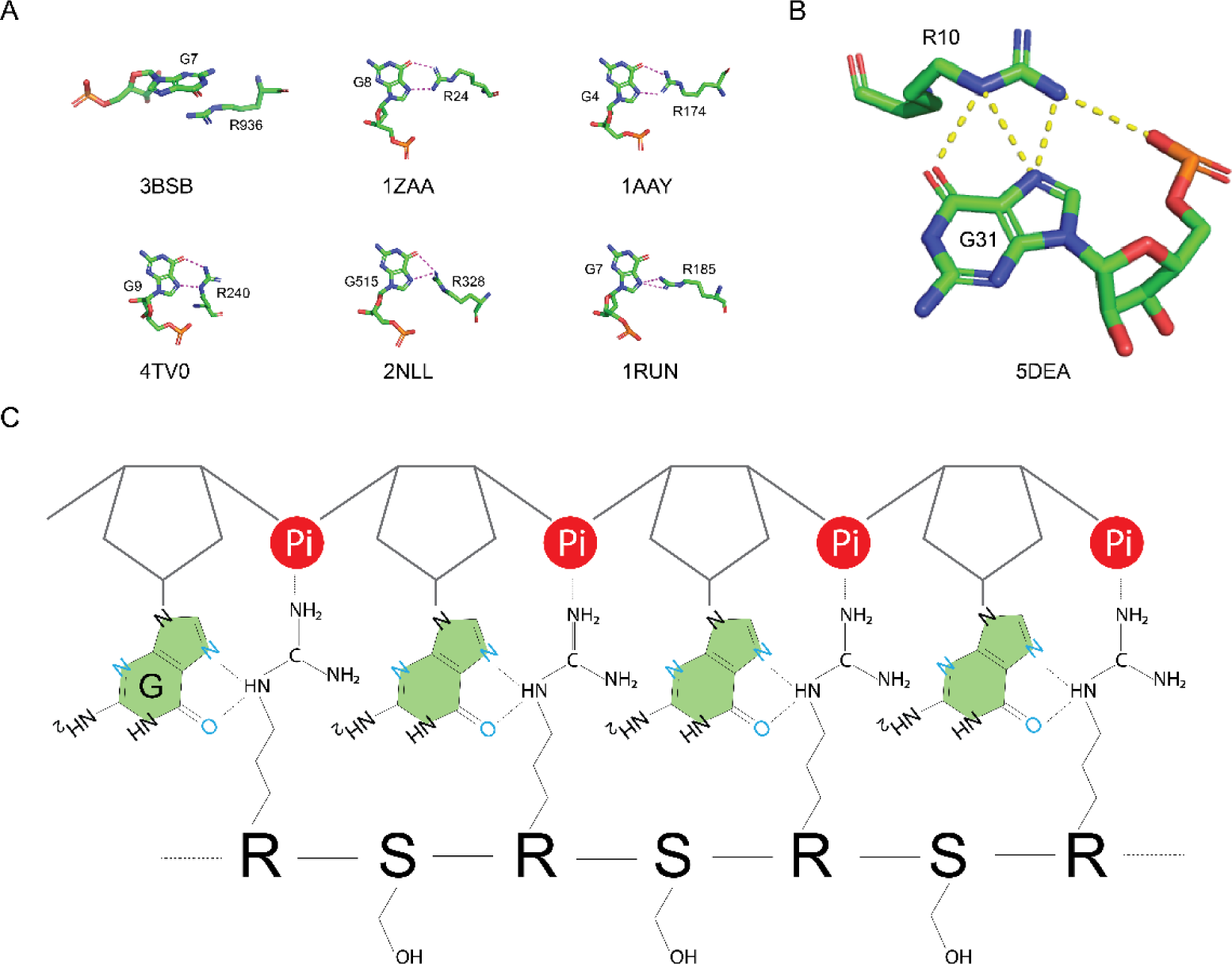
Proposed mechanism by which RS binds unfolded GQ-forming sequences. (A) Example stacking interaction or H-bonds between Arg and guanine, with each example accompanied by its residue number and four-letter PDB ID. Dotted lines represent H-bonds. (B) A representative interaction is depicted in which the Arg sidechain simultaneously forms H-bonds and a salt bridge with the guanine base and phosphate backbone. (C) A schematic representation of the interaction between RS and a G-rich RNA region. In the extended conformation, the distance of neighboring of nucleotides (6.5-7.5 Å) matches that of neighboring the Arg-Ser dipeptide (6.8 Å).

In a recent genome-wide bioinformatics study, a notable enrichment of GQ sequences has been observed within the exon/intron boundaries(51). It is well-established that these boundaries serve as a critical platform for recruiting numerous splicing factors and spliceosome components. Consequently, it is plausible that GQ structures represent an additional layer of regulation influencing both constitutive and alternative splicing events. For quite some time, the impact of RNA secondary structure on alternative splicing has been acknowledged, with SR proteins emerging as potential regulators in this context. However, the precise mechanisms governing the collaborative influence of SR proteins and GQ RNA on the splicing process remain a subject that needs further in-depth investigation and exploration.

## Data Availability

The data underlying this article are available in the article and in its online supplementary material.

## Acknowledgements

We want to thank UAB Central Alabama High-Field NMR Facility. This work is supported by U.S. National Institutes of Health, NIGMS.

