## Supplemental data_V3 for "Unearthing SRSF1’s Novel Function in Binding and Unfolding of RNA G-Quadruplexes"

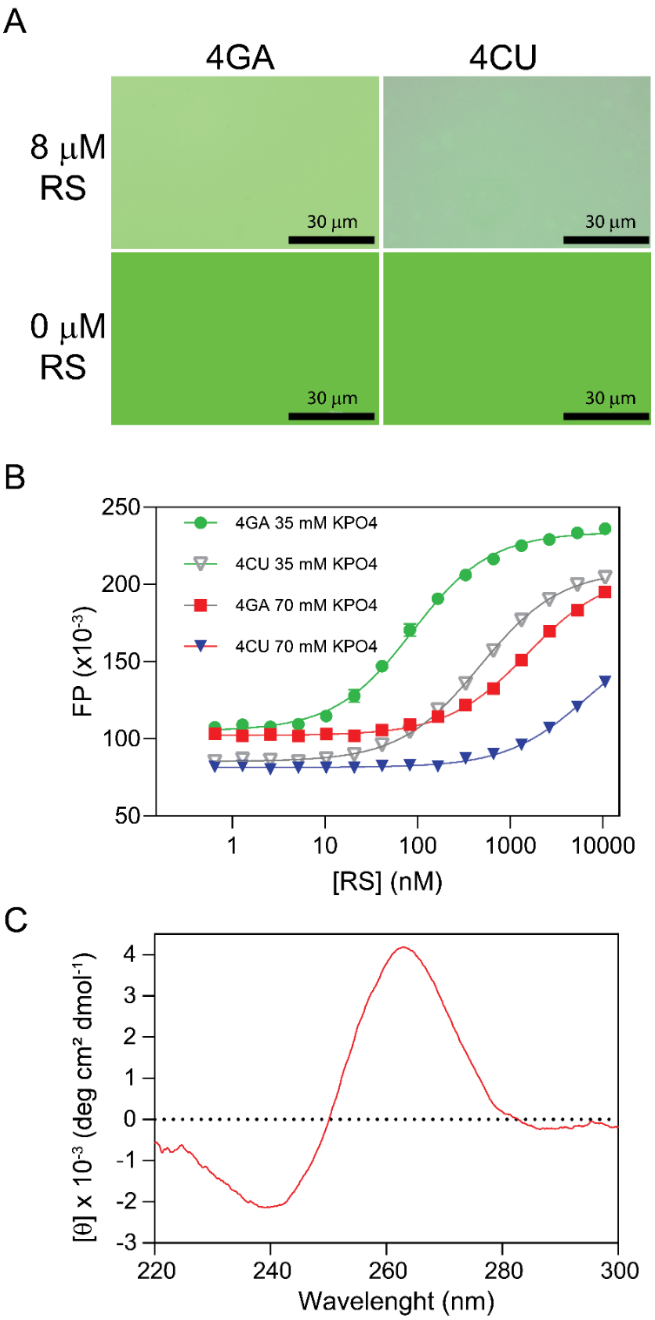


Figure S1: (A) RS does not form phase separation with 8-mer RNA in the FP binding buffer that contains 35 mM $K_3PO_4$, 2.5 mM NaCl, 15 mM HEPES pH 7.5, 0.015% TWEEN 20, and 0.075 mM TCEP. The images were recorded by a Cytation 5 plate reader. The blank bars indicate 30 µm. (B) Fluorescence polarization binding profiles for RS with 4GA and 4CU nucleotides in 35 and 70 mM $K_3PO_4$. RNA oligomers were labeled by fluorescein at the 5’ end, and the fluorescent probe concentration was 10 nM for all binding assays. (C) Circular spectrum of 8G RNA. The spectrum was collected with 5 µM of RNA in 35 mM $K_3PO_4$, 2.5 mM NaCl, 15 mM HEPES pH 7.5, 0.015% TWEEN 20, 0.075 mM TCEP.

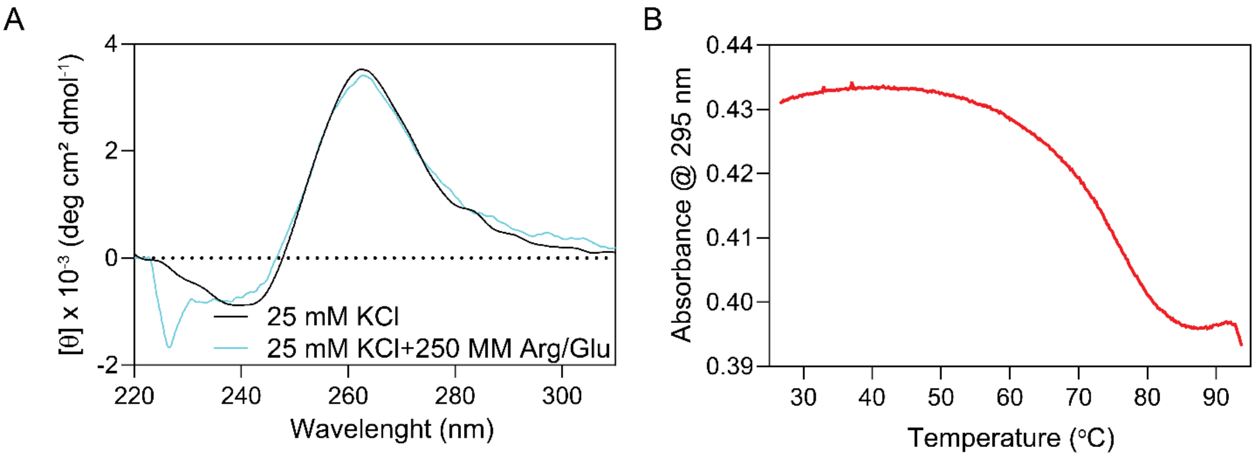


Figure S2: The ARPC2 GQ RNA sequence forms G-quadruplex. (A) Circular dichroism spectra of ARPC2 GQ in 25 mM KCl with and without 250 mM Arg/Glu. Addition of Arg/Glu does not change the secondary structure of ARPC2 GQ. Due to strong absorbance of Arg/Glu, the CD spectrum region lower than 230 nm is not usable. (B) The UV melting curve of ARPC2 GQ. The data was collected at a concentration of 5 μM in a buffer containing 25 mM KCl, 0.25 M Arg/Glu, 20 mM Tris-HCl, pH 7.5, 0.1 mM EDTA, and 0.1 mM TCEP.

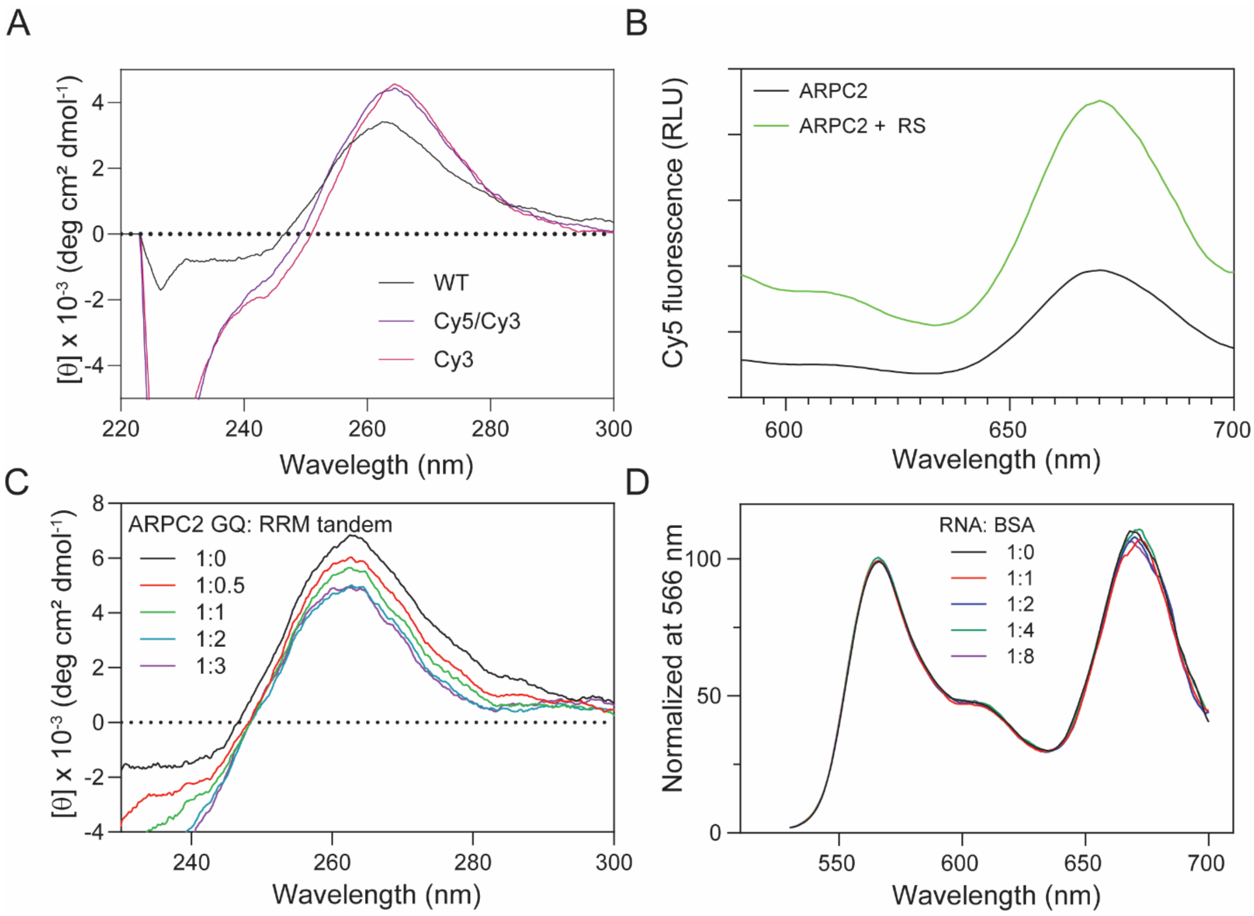


Figure S3: (A) Fluorophore labeling does not affect the ARPC2 GQ structure. CD spectra were collected at 5 µM. (B) The Cy5 fluorescence of Cy5-ARPC2 GQ-Cy3 with and without 400 nM RS. The RNA concentration is apo RNA is 200 nM. The spectrum with RS was normalized to the same concentration as the apo RNA. The samples were excited with a light at 500 nm. The Y-axis is in the relative light unit (RLU). (C) Circular dichroism of 5 µM of ARPC2 GQ titrated with different concentration of the RRM tandem. (D) Fluorescence spectra of 200 nM ARPC2 GQ titrated with different concentration of BSA. The spectra were recorded with exciting light at 500 nm. All fluorescence and CD spectra were collected in 25 mM KCl, 0.25 M Arg/Glu, 20 mM Tris-HCl pH 7.5, 0.1 mM EDTA, 0.1 mM TCEP, and 0.02% TWEEN 20.

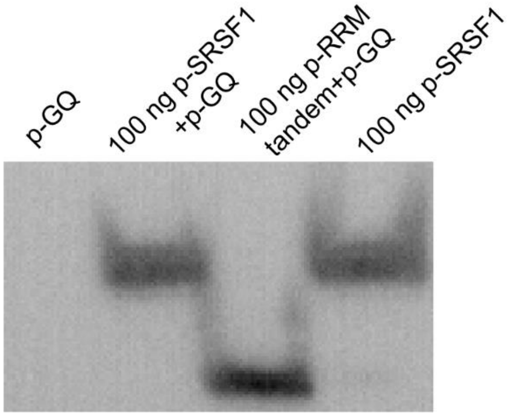


Figure S4: Confirmation of SRSF1 and the RRM tandem expression by western blot analysis.
